# NanoPack: visualizing and processing long read sequencing data

**DOI:** 10.1101/237180

**Authors:** Wouter De Coster, Svenn D’Hert, Darrin T. Schultz, Marc Cruts, Christine Van Broeckhoven

**Author notes:** To whom correspondence should be addressed Wouter de Coster, MSc Neurodegenerative Brain Diseases research group, VIB Center for Molecular Neurology, University of Antwerp – CDE, Universiteitsplein 1, 2610 Antwerp, Belgium.

## Abstract

**Summary:** Here we describe NanoPack, a set of tools developed for visualization and processing of long read sequencing data from Oxford Nanopore Technologies and Pacific Biosciences.

**Availability and Implementation:** The NanoPack tools are written in Python3 and released under the GNU GPL3.0 Licence. The source code can be found at https://github.com/wdecoster/nanopack, together with links to separate scripts and their documentation. The scripts are compatible with Linux, Mac OS and the MS Windows 10 subsystem for linux and are available as a graphical user interface, a web service at http://nanoplot.bioinf.be and command line tools.

**Supplementary information:** Supplementary tables and figures are available at Bioinformatics online.

## Introduction

The dominant sequencing by synthesis technology is characterized by sequencing a fixed short read length template (50-300bp) with high accuracy (error rate <1%) (Goodwin, McPherson, and McCombie 2016). In contrast, long read sequencing methods from Oxford Nanopore Technologies (ONT) and Pacific Biosciences routinely achieve read lengths of 10kb, with a long tail of up to 1.2 Megabases for ONT (unpublished results). These long reads come with a tradeoff of lower accuracy of about 85-95% (Jain, Tyson, et al. 2017; Giordano et al. 2017). It is evident that these characteristics make many existing Illumina-tailored QC tools, such as FastQC (Babraham Bioinformatics 2010), suboptimal for long read technologies. NanoPack, a set of Python scripts for visualizing and processing long read sequencing data, was developed to partially bridge this gap. Earlier tools such as poretools (Loman and Quinlan 2014), poRe (Watson et al. 2015) and IONiseR (Smith 2017) mainly focussed on feature extraction from the older fast5 file formats, and alternative tools such as pycoQC (aleger 2017) and minion_qc (roblanf n.d.) do not offer the same flexibility and options as NanoPack. The plotting style from the pauvre tool (Schultz n.d.) got incorporated in NanoPack (supplementary figure S3).

## Software description

### Installation and dependencies

NanoPack and individual scripts are available through the public software repositories PyPI using pip and bioconda through conda (Dale et al. 2017). The scripts build on a number of third party Python modules: matplotlib (Hunter 2007), pysam (Li et al. 2009; Heger 2009), pandas (McKinney 2011), numpy (Walt, Colbert, and Varoquaux 2011), seaborn (Waskom et al. 2017) and biopython (Cock et al. 2009).

### Scripts for statistic evaluation and visualization

NanoStat produces a comprehensive statistical data summary (Supplementary table 2). NanoPlot and NanoComp produce informative QC graphs displaying multiple aspects of sequencing data (Figure 1, Supplementary table 1) and accept input data in (compressed) fastq or fasta format, bam and (compressed) albacore summary files or multiple files of the same type. All plots and summary statistics are combined in an html report. Because long and variable read lengths may be challenging to interpret on a linear axis, there is also an option to plot the read lengths on a log scale. Plots can be produced in standard image file formats including png, jpg, pdf and svg. NanoPlot produces read length histograms, cumulative yield plots, violin plots of read length and quality over time, and bivariate plots comparing the relationship between read lengths, quality scores, reference identity and read mapping quality. Better insight in big datasets can be obtained using bivariate plots with a 2D kernel density estimation or hexagonal bins (Figure 1E, F, S3). Optional arguments include random downsampling of reads and removing all reads above a length cutoff or below a quality cutoff. Data from a multiplexed experiment in albacore summary format can be separated, resulting in plots and statistics per barcode. NanoComp performs comparison across barcodes or experiments of read length and quality distributions, number of reads, throughput and reference identity.

**Figure 1.**
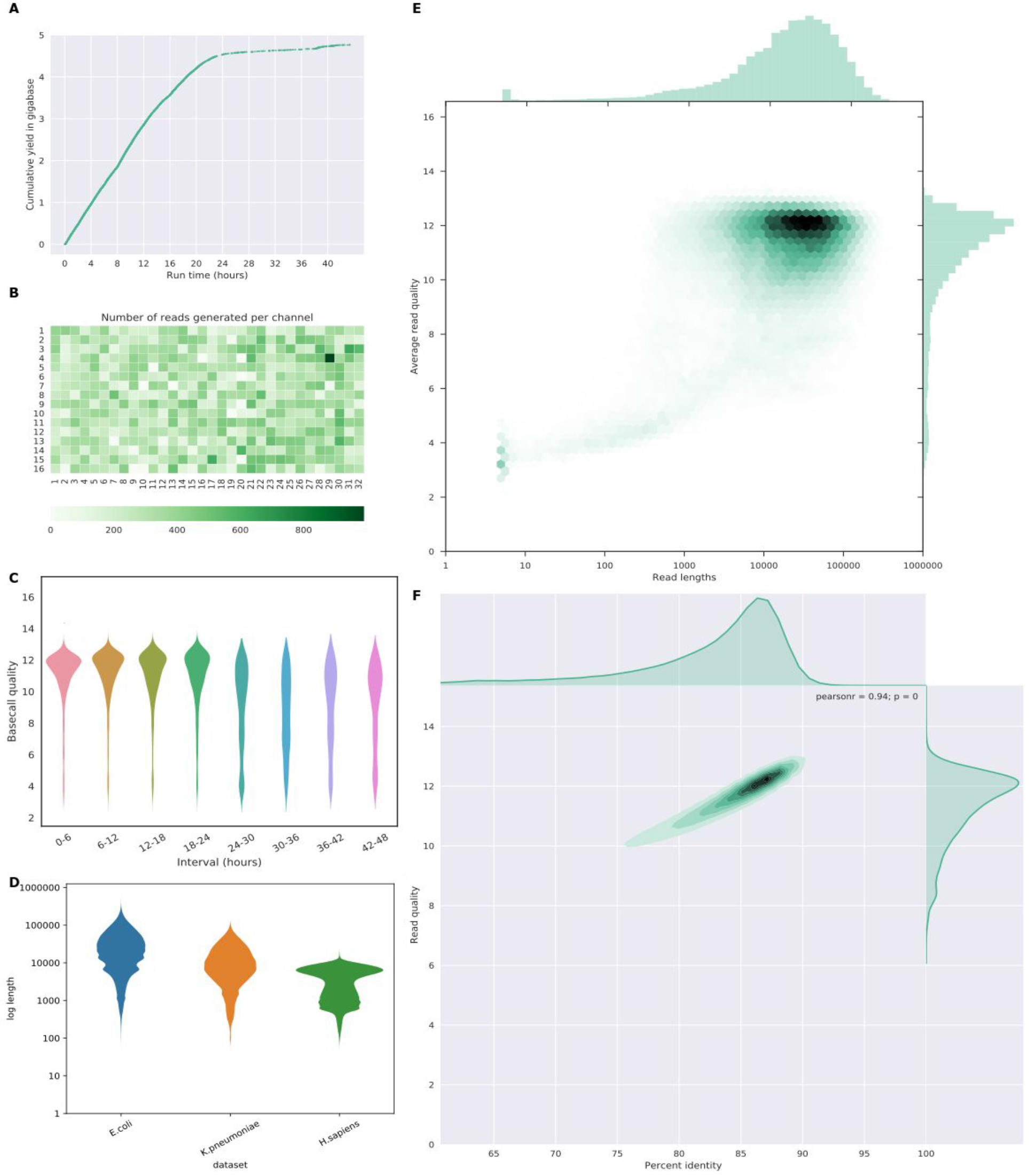
Examples of plots of NanoPlot and NanoComp. **A** Cumulative yield plot **B** Flow cell activity heatmap showing number of reads per channel. **C** Violin plots comparing base call quality overtime. **D** NanoComp plot comparing log transformed read lengths of the E.coli dataset with a K.pneumoniae and human dataset. **E** bivariate plot of log transformed read length against base call quality with hexagonal bins and marginal histograms. **F** bivariate plot of base call quality against percent identity with a kernel density estimate and marginal density plots.

### Scripts for data processing

NanoFilt and NanoLyse were developed for processing reads in streaming applications and therefore have a minimal memory footprint and can be integrated in existing pipelines prior to alignment. NanoFilt is a tool for read filtering and trimming. Filtering can be performed based on mean read quality, read length and mean GC content. Trimming can be done with a user-specified number of nucleotides from either read ends. NanoLyse is a tool for rapid removal of contaminant DNA, using the Minimap2 aligner through the mappy Python binding (Li 2017). A typical application would be the removal of the lambda phage control DNA fragment supplied by ONT, for which the reference sequence is included in the package. However, this approach may lead to unwanted loss of reads from regions highly homologous to the lambda phage genome.

## Examples and discussion

The NanoPlot and NanoComp examples (Figure 1) are based on an ONT E.coli dataset from an ultra-long read protocol sequenced on an R9.4 MinION flow cell (Quick and Loman 2017) generating 150,735 reads, base called using Albacore 2.0.2 and aligned to the E.coli reference genome using Minimap2 (Li 2017). The cumulative yield (figure 1A) shows a lower efficiency when the flow cell wears out. A heat map of the physical layout of the MinION flow cell (Figure 1B) highlights more productive channels and could potentially identifying suboptimal loading conditions, such as introduction of an air bubble. The mean base call quality per 6 hours interval (Figure 1C) shows a uniform high quality in the beginning, with lower quality reads after 24h. In a bivariate plot comparing log transformed read lengths with their mean quality score (Figure 1E) the majority of reads can be identified at lengths of 10kb and quality scores of 12 by the color intensity of the hexagonal bins, with a subgroup of low quality short reads. Plotting the mean quality against the per read percent reference identity (as a proxy for accuracy) (Figure 1F) highlights a strong correlation, here with the number of reads plotted using a kernel density estimate. Additional examples from NanoPlot can be found in the supplementary information online, including standard and log transformed histograms, optionally with the N50 metric (S1,2) and a bivariate plot comparing effective read length with aligned read length (S4), identifying reads which are only partially aligned to the reference genome.

The NanoComp plot (figure 1D) compares the log transformed read lengths of the same E.coli dataset to a K. pneumoniae (Wick et al. 2017) and a human PromethION dataset (unpublished), clearly showing differences in the length profile with far longer reads in the E.coli dataset, standard read lengths in the library prep by ligation from K. pneumoniae and suboptimal read lengths from the human sample. Additional examples from NanoComp can be found in the supplementary information online, indicating that the K. pneumoniae library has both the highest yield (S5) and on average higher quality scores (S6) than both the human and E.coli dataset, but a comparable percent identity (S7) with the human dataset.

## Conclusion

NanoPack is a package of efficient Python scripts for visualization and processing of long read sequencing data available on all major operating systems. Installation from the PyPI and bioconda public repositories is trivial, automatically taking care of dependencies. The plotting tools are flexible and customizable to the users need. Using a single NanoPlot or NanoComp command a full html report containing all summary statistics and plots can be prepared, and the software is easily accessible through the graphical user interface and web service, in addition to the command line scripts.

## Acknowledgments

The authors acknowledge Mick Watson for contributing the mean GC content filtering to NanoFilt and Andreas Sjödin for maintaining the bioconda build recipes. The authors are also thankful to the many users who provided helpful suggestions and feature requests for these scripts.

